# Pharmacometabolomics reveals urinary diacetylspermine as a biomarker of doxorubicin effectiveness in triple negative breast cancer

**DOI:** 10.1101/2022.01.19.475906

**Authors:** Thomas J. Velenosi, Kristopher W. Krausz, Keisuke Hamada, Tiffany H. Dorsey, Stefan Ambs, Shogo Takahashi, Frank J. Gonzalez

**Author notes:** **Correspondence:** Frank J. Gonzalez, Ph.D., Chief, Laboratory of Metabolism, National Cancer Institiute, National Institutes of Health, 37 Convent Drive, Room 3106, Bethesda, MD 20892, Thomas J. Velenosi, Ph.D., Assitant Professor, Faculty of Pharmaceutical Sciences, University of British Columbia, 2495 Wesbrook Mall, Vancouver, BC, Canada, V6T 1Z3. Senior author (Shogo Takahashi).

## Abstract

Triple-negative breast cancer (TNBC) patients receive chemotherapy treatment, including doxorubicin, due to the lack of targeted therapies. Drug resistance is a major cause of treatment failure in TNBC and therefore, there is a need to identify biomarkers that determine effective drug response. Here, a pharmacometabolomics study was performed using TNBC patient-derived xenograft models to detect urinary metabolic biomarkers of doxorubicin effectiveness. Diacetylspermine was identified as a urine metabolite that robustly changed in response to effective doxorubicin treatment, which persisted after the final dose. Diacetylspermine was directly traced back to the tumor and correlated with tumor volume. Ex vivo tumor slices revealed that doxorubicin directly increases diacetylspermine production by increasing tumor spermidine/spermine N^1^-acetyltransferase 1 expression and activity, which was corroborated by elevated polyamine flux. In breast cancer patients, tumor diacetylspermine was elevated compared to matched non-cancerous tissue and increased in HER2+ and TNBC compared to ER+ subtypes. In addition, 12-hour urine diacetylspermine was associated with breast cancer tumor volume and poor tumor grade. This study describes a pharmacometabolomics strategy for identifying cancer metabolic biomarkers that indicate drug response. Our findings characterize urine diacetylspermine as a non-invasive biomarker of doxorubicin effectiveness in TNBC.

## Introduction

Women have a 1-in-8 chance of developing breast cancer, which is the second leading cause of cancer-related death (1). Triple negative breast cancer accounts for 10-20% of breast cancers (2). In the absence of targets to exploit with therapy, TNBC patients are treated with aggressive chemotherapy as the primary systemic treatment. TNBC chemotherapy regimens typically contain a combination of taxanes, cyclophosphamide, and anthracyclines such as doxorubicin, to maximize therapeutic response. Only 30-40% of patients with TNBC who receive taxane- and anthracycline-based therapy will achieve a pathological complete response (3). TNBC has the worst prognosis of all breast cancer subtypes, with the highest 5-year mortality across all disease stages (4). Recurrence is frequent in TNBC patients with most events occurring within 3 years of the disease diagnosis (5). Current techniques for detecting TNBC recurrence rely on imaging and are limited by many factors including tumor size, individual breast characteristics, skill of the examiner, and follow-up to minimize false negatives (6). Consequently, a recurrence can only be detected months after primary therapy. Therefore, identification of clinical biomarkers that can be used to detect a drug response during treatment would provide a clinical benefit to patients with TNBC.

Clinical biomarkers that rely on gene mutations and expression patterns are obtained through biopsy sampling, and thus frequent assessment is impractical. Metabolites can provide a reliable prognostic and diagnostic readout for disease status (7). By extending the use of metabolic biomarkers to pharmacometabolomics, metabolites excreted from tumors can be measured by routine non-invasive sampling before and throughout the duration of drug treatment to monitor therapeutic effectiveness. However, factors including BMI, age and sex can greatly influence metabolic biomarkers (8). When combined with the heterogeneity of TNBC tumors, it can be extremely challenging to identify tumor and treatment specific metabolic perturbations. Indeed, clinical metabolomics studies require many subjects to be properly powered and lack the ability to identify the source of biomarker production. Therefore, an innovative approach is needed to identify and characterize pharmacometabolomic biomarkers of cancer treatment.

Cell lines and cell-line xenografts have historically served as the foundations of preclinical cancer research. These models have been used to identify metabolic biomarkers specific to breast cancer and drug metabolic response (9-11). However, cultured cell lines adapt *in vitro* and genetically diverge from the original tumors (12). Moreover, breast cancer cell lines do not sufficiently capture the tumor microenvironment and heterogenous cell population (13). For these reasons, *in vitro* and xenograft models using cell lines diverge from the tumor cells in breast cancer patients, and often fail to predict clinical success. The recent development of patient-derived xenografts (PDX), which can be grown and passaged for multiple generations while retaining their genetic and molecular integrity, provide a well-suited model to test clinically relevant hypotheses.

In this study, we employed TNBC-PDX models known to be sensitive or resistant to doxorubicin as an innovative pharmacometabolomics approach to identify and characterize urinary metabolic biomarkers of doxorubicin effectiveness. With this approach, we identified diacetylspermine, a catabolic product in the polyamine pathway, as a urinary metabolic biomarker that increases with increasing tumor size, and can be traced back to the implanted tumor. Using tumor ex vivo tissue slices and applying stable isotope tracer substrates for the polyamine pathway, we further show that the effect of doxorubicin on SAT1 expression and its activity is the mechanism of increased diacetylspermine. Finally, these observations were extended to human breast cancer by showing that urinary diacetylspermine levels are associated with tumor size in a large clinical study and that tumor diacetylspermine is elevated in tumor tissue compared to adjacent noncancerous tissue.

## Results

### Diacetylspermine is a metabolic biomarker of doxorubicin effectiveness

To identify metabolic biomarkers of doxorubicin effectiveness, we treated TM97 (doxorubicin-resistant) and TM98 (doxorubicin-sensitive) TNBC-PDX mice with vehicle or doxorubicin and collected urine for 24 hours after each dose. A final urine collection was obtained in the absence of drug treatment to limit drug metabolite interference during metabolomics analysis. Doxorubicin treatment reduced tumor growth in the TM98 TNBC-PDX model, but had no effect on TM97, confirming sensitivity and resistance, respectively (Figure 1B). Principal component analysis using the metabolome data from day 21 urine samples showed separation of tumor-bearing animals by doxorubicin treatment in TM98, which was partially explained by accumulation of one metabolite, diacetylspermine (Figure 1C,D). Univariate statistical analysis revealed diacetylspermine as the most significantly altered urine metabolite as a product of the interaction of tumor & treatment and time (Figure 1E, Supplemental Table 1). Urine diacetylspermine levels were significantly increased in mice implanted with TM98 when compared to TM97. Interestingly, doxorubicin significantly increased TM98 urinary diacetylspermine levels, despite the corresponding decrease in tumor size during treatment. When quantified, urine diacetylspermine increased 8.5-fold and 12.8-fold on day 1 and on day 21, respectively, in TM98 mice after treatment with doxorubicin and compared to the vehicle control group (Figure 1F). In TM97, urine diacetylspermine was not significantly altered throughout tumor growth and regardless of treatment. Tumor volume was correlated with urine diacetylspermine with a particularly strong correlation for TM98 control treated mice (r^2^ = 0.8, *P* < 0.001, Figure 1G).

**Figure 1.**
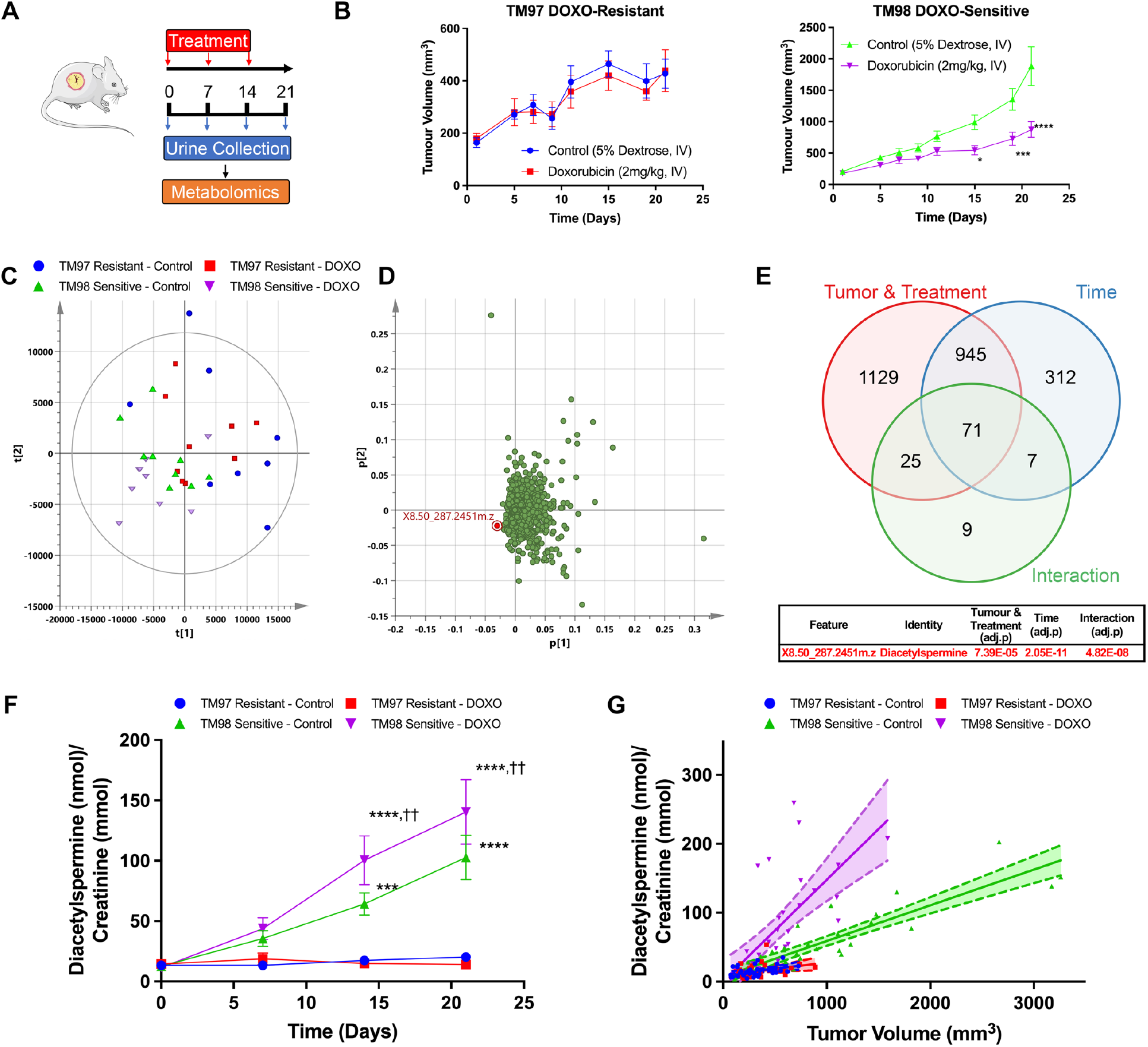
Longitudinal urine metabolomics identifies diacetylspermine as a metabolic biomarker of doxorubicin effectiveness. (**A**) Experimental design for urine collection of TNBC-PDX TM97 and TM98 mice treated with vehicle (5% dextrose, IV) or doxorubicin (DOXO, 2 mg/kg, IV). (**B**) Tumor growth curves showing TM97 resistance and TM98 sensitivity to DOXO treatment.(**C**) Principal component analysis and loadings plot of untargeted urine metabolomics on day 21.(**D**) Venn diagram showing number of significantly altered features with X8.56_287.2451m.z. being the most significant by two-way repeated measures ANOVA. (**E**) Quantified urinary diacetylspermine and **(F)** correlation with tumor volume. Data are presented as mean ± s.e.m. or individual values, \*\*\**P*<0.001, \*\*\*\**P*<0.0001 vs TM97, ^††^*P*<0.01 vs TM98 control; n=8-9. Significance was determined by one-way ANOVA with Tukey’s test or two-way repeated measures ANOVA with Sidak’s correction. Correlation plots display 95% confidence intervals.

### Urine diacetylspermine is increased by tumor generated diacetylspermine through doxorubicin-mediated induction of SAT1

The correlation between tumor volume and urinary diacetylspermine that we detected suggested that diacetylspermine may be directly produced by the implanted TNBC-PDX. Therefore, we hypothesized that increased urinary diacetylspermine levels could be a result of tumor SAT1 induction, as SAT1 is commonly upregulated in a variety of cancers including breast cancer. Plasma diacetylspermine was significantly increased in doxorubicin treated TM98 when compared to TM97. Yet, plasma levels were generally low and near our limit of quantification (LOQ, Figure 2A). Tumor levels of other spermine/spermidine-related metabolites like N1-acetylspermidine, N-acetylspermine and diacetylspermine, as well as SAT1 expression, were significantly elevated in TM98 compared with TM97 mice (Figure 2D,E). Notably, the levels of these metabolites were further increased by doxorubicin treatment in the TM98 doxorubicin responder mice, but not in TM97 mice.

**Figure 2.**
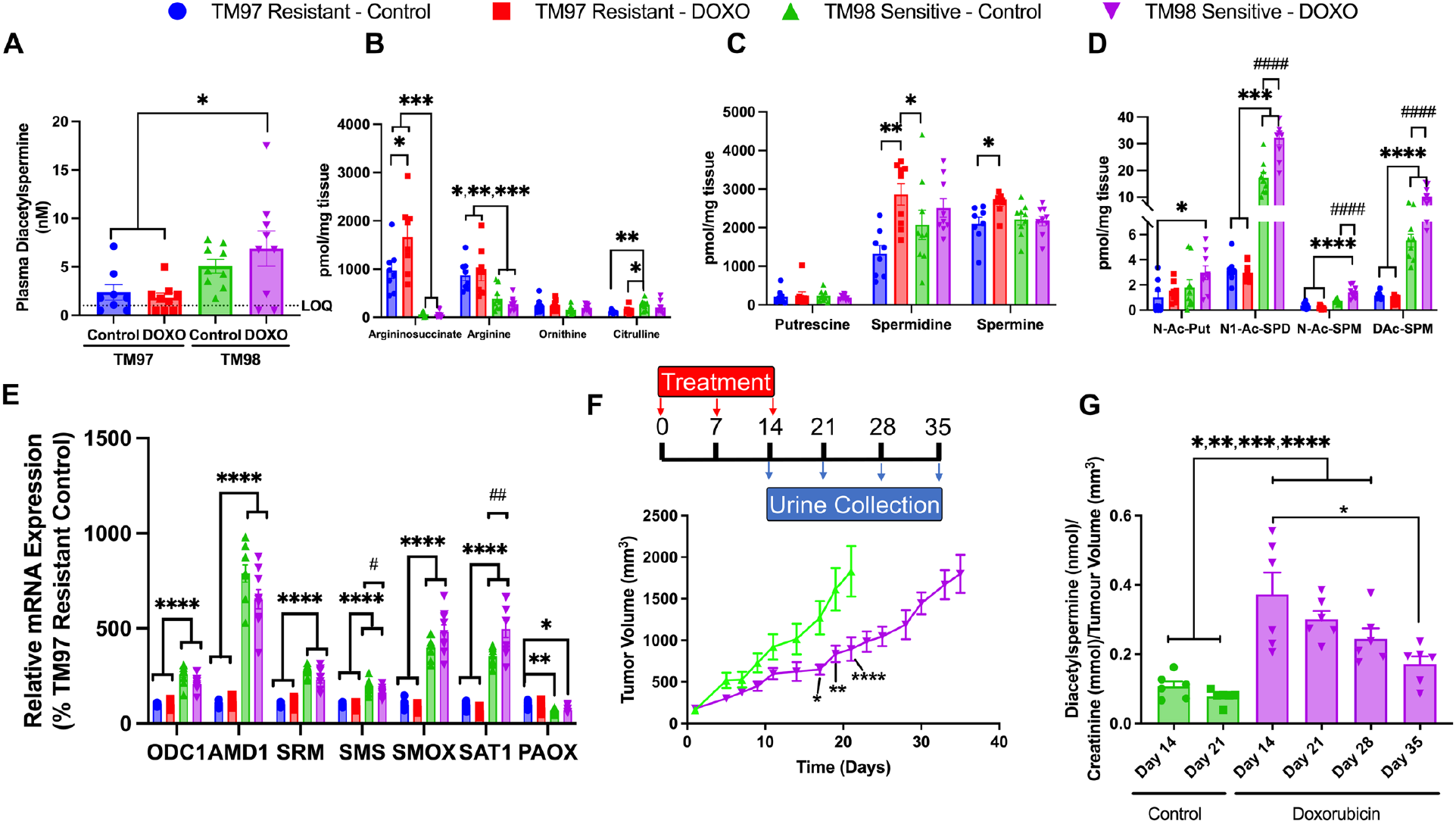
Doxorubicin treatment increases SAT1 expression and tumor production of diacetylspermine, which is sustained following the final dose. (**A**) Plasma diacetylspermine and tissue urea cycle (**B**), polyamine (**C**) and acetylated polyamine (**D**) metabolites from TM97 and TM98 TNBC-PDX mice treated with vehicle (5% dextrose, IV) or doxorubicin (DOXO, 2 mg/kg, IV). (**E**) mRNA expression of polyamine pathway enzymes in TM97 and TM98 TNBC-PDX mice treated with vehicle or doxorubicin. (**F**) TM98 mice treated with vehicle or doxorubicin and placed in metabolic cages for urine collection as indicated. (**G**) Urine diacetylspermine normalized to tumor volume. Data are presented as mean ± s.e.m., n = 8-9 (**A-E**), n = 6 (**H-G**), \**P*<0.05, \*\**P*<0.01, ***P<0.01, \*\*\*\**P*<0.0001 vs TM97; # *P*<0.05, ##*P*<0.01, ####*P*<0.0001 vs TM98 control. Plasma concentrations below limit of quantification (LOQ) were calculated as LOQ/2. Significance was determined by one-way ANOVA with Tukey’s test or two-way repeated measures ANOVA with Sidak’s correction.

### Increased SAT1 expression and function by doxorubicin is time and dose-dependent

To confirm whether doxorubicin directly induced SAT1, we treated TNBC cell lines with physiologically relevant levels of doxorubicin (1 μM) over a time course. Significant induction of SAT1 by doxorubicin did not occur until 24 hours after treatment, which was consistent across cancer cell lines derived from various tumor types (Supplemental Figure 1).

TM98 mice given doxorubicin continued to produce elevated urine diacetylspermine and upregulated SAT1 expression when compared to control mice at one week after the final dose. To further characterize the pharmacodynamic response, mice given doxorubicin were placed into metabolic cages weekly from day 14 to 35 (Figure 2F). When normalized to tumor volume, the increase in urine diacetylspermine was sustained for 14 days following the final dose. Therefore, doxorubicin caused a persistent increase in diacetylspermine production long after the drug was cleared (Figure 2G).

Next, we determined if doxorubicin affected the level of SAT1 induction in a dose-dependent manner. Doxorubicin dose-dependently increased SAT1 expression similarly in TNBC cells and TM98 ex vivo tissue slices with the most robust increase occurring at 1 μM (Supplemental Figure 2A). Furthermore, elevated acetylated polyamine levels coincided with the increase in SAT1 expression (Supplemental Figure 2B). Together, these results indicate that elevated urinary diacetylspermine is the result of increased SAT1 expression.

### TNBC ex vivo slices recapitulate in vivo SAT1 induction by doxorubicin

To further characterize the effect of doxorubicin on the polyamine pathway, we incubated TM97 and TM98 ex vivo tissue slices with doxorubicin, or vehicle control. After 24-hour doxorubicin treatment, acetylated polyamines were significantly increased in TM98 compared to control or compared to TM97 ex vivo slices (Figure 3C). This occured without affecting tissue slice viability (Supplemental Figure 2C). Although metabolites were measured 1 and 7 days *after ex vivo* slice and in vivo treatment, respectively, *ex vivo* slices largely captured the in vivo urea cycle and polyamine metabolic profile (Figure 2B-D and Figure 3A-C). One noted difference was the increase in putrescine following 24-hour doxorubicin treatment suggesting an early onset response to polyamine depletion, which may dissipate in the days following in vivo treatment (Figure 2C and Figure 3B).

**Figure 3.**
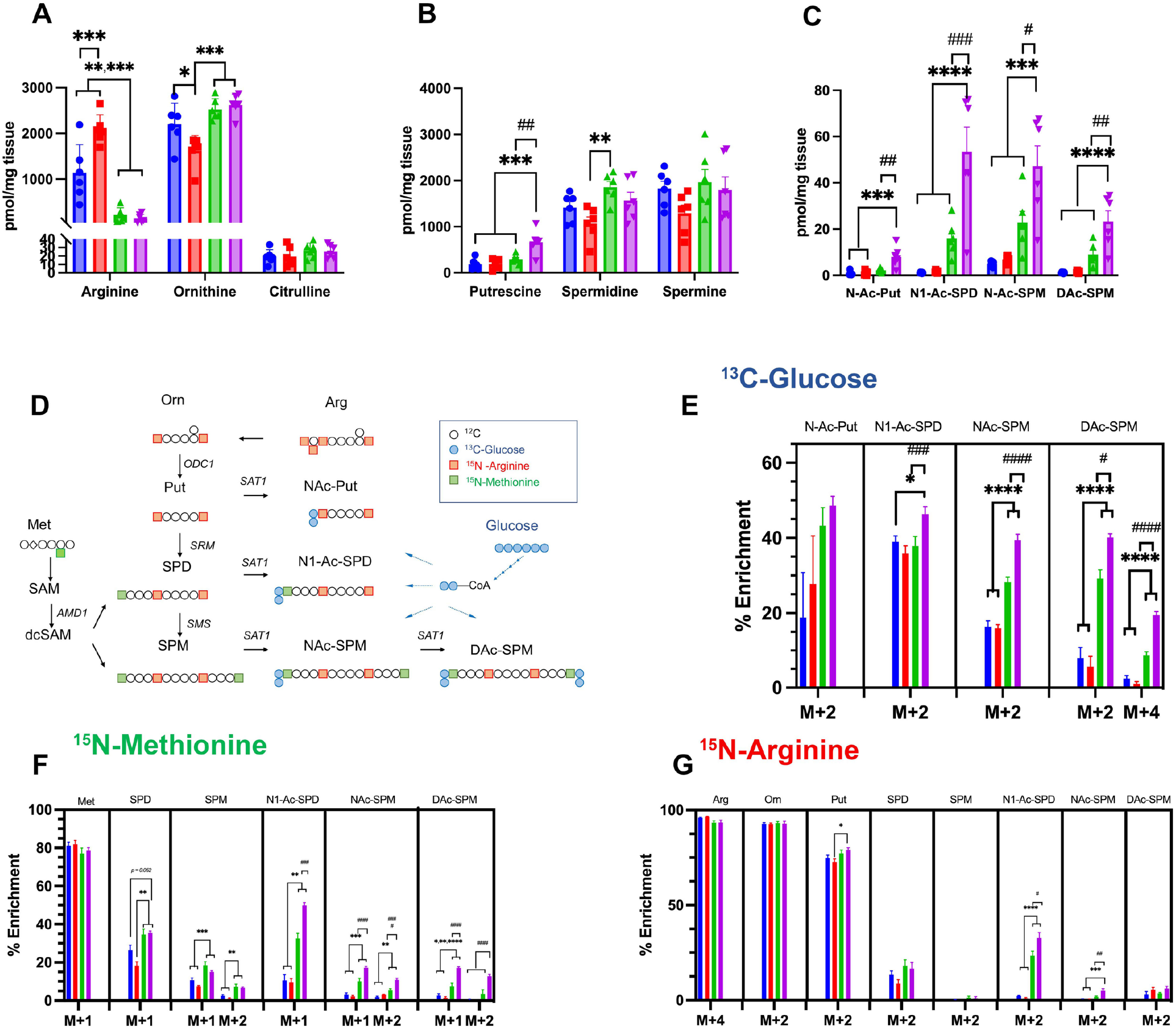
S-Adenosylmethioninamine and acetylated polyamine flux contribute to diacetylspermine production in *ex vivo* TNBC-PDX tissue slices. Quantified urea cycle (**A**), polyamine (**B**) and acetylated polyamine (**C**) metabolites. (**D)** atom transition map of ^13^C-glucose (**E**), ^15^N-methionine (**F**), and ^15^N-arginine (**G**) stable isotope metabolic flux in TM97 and TM98 ex vivo tissue slices treated with vehicle (DMSO) or 1 μM doxorubicin for 24 hours. Experiments were performed in triplicate. Data are presented as mean ± s.e.m., n = 6. \**P*<0.05, \*\**P*<0.01, \*\*\**P*<0.001, \*\*\*\**P*<0.0001 vs TM97; ^#^ *P*<0.05, ^##^ *P*<0.01, ^###^ *P*<0.001, ^####^ *P*<0.0001 vs TM98 control. Significance was determined by two-sided one-way ANOVA with Tukey’s test.

### Polyamine flux contributes to TNBC diacetylspermine production

Besides SAT1, the rate limiting polyamine anabolic enzymes, ornithine decarboxylase 1 (ODC1) and adenosylmethionine Decarboxylase 1 (AMD1), were also increased in TM98 compared to TM97 (Figure 2E), however, polyamines levels were largely unaltered (Figure 1C). Therefore, we hypothesized that increased diacetylspermine tumor production is the result of increased metabolic flux through the urea cycle and polyamine pathways. To delineate these pathways, we incubated TM97 and TM98 *ex vivo* tissue slices with media containing pathway-selective stable isotope-labelled substrates. TM98 *ex vivo* slices cultured in the presence of ^13^C-glucose showed a significantly elevated incorporation of the ^13^C-label into acetylated spermine metabolites compared to TM97. Moreover, the magnitude of increase was greatest for diacetylspermine (Figure 3E). The postulated increased polyamine flux in TM98, when compared to TM97, was further verified by ^15^N-methionine enrichment of spermidine and spermine (Figure 3F), while total metabolite levels remained mostly unaltered (Figure 3B). A substantially larger fraction of N1-acetylspermidine and N-acetylspermine were +2 labelled suggesting elevated flux of newly generated polyamines towards acetylation in TM98 (Figure 3F). This flux was further increased by doxorubicin treatment. During ^15^N-arginine treatment, labeled putrescine accumulated and the stable isotope incorporation pattern was similar to other isotopes for N1-acetylspermidine and N-acetylspermine (Figure 3G). However, minimal label incorporation occurred in spermidine and spermine.

Polyamines are highly regulated and, when reaching high levels, synthesis is quickly reduced while catabolism increases. To test whether the polyamine pathway was capable of responding to elevated polyamine levels in TNBC-PDX, we treated TNBC-PDX *ex vivo* slices with vehicle or doxorubicin in the presence of spermine. Spermine treatment dose-dependently decreased putrescine and acetylated polyamines levels in control and doxorubicin treated TM98 *ex vivo* tissue slices (Supplemental Figure 3). Spermidine and spermine levels remained unaffected but, at high spermine concentrations, polyamine regulation failed leading to increased diacetylspermine in TM97 and TM98, thus confirming that SAT1 can respond to elevated polyamines to generate diacetylspermine in both tumors (Supplemental Figure 3G).

### Doxorubicin decreases urinary diacetylspermine in TNBC with high baseline SAT1 function

To further validate that diacetylspermine is a metabolic biomarker of doxorubicin drug response, we used two additional PDX models and treated TM89 (doxorubicin-resistant) and TM99 (doxorubicin-sensitive) TNBC-PDX mice with doxorubicin or vehicle. Tumor growth rates were similar between TM89 and TM99 (Figure 4A). Urine diacetylspermine levels were increased in TM99 compared to TM89, which correlated with tumor volume and changed similarly in plasma, tumor tissue and *ex vivo* tumor slices (Figure 4B, C).

**Figure 4.**
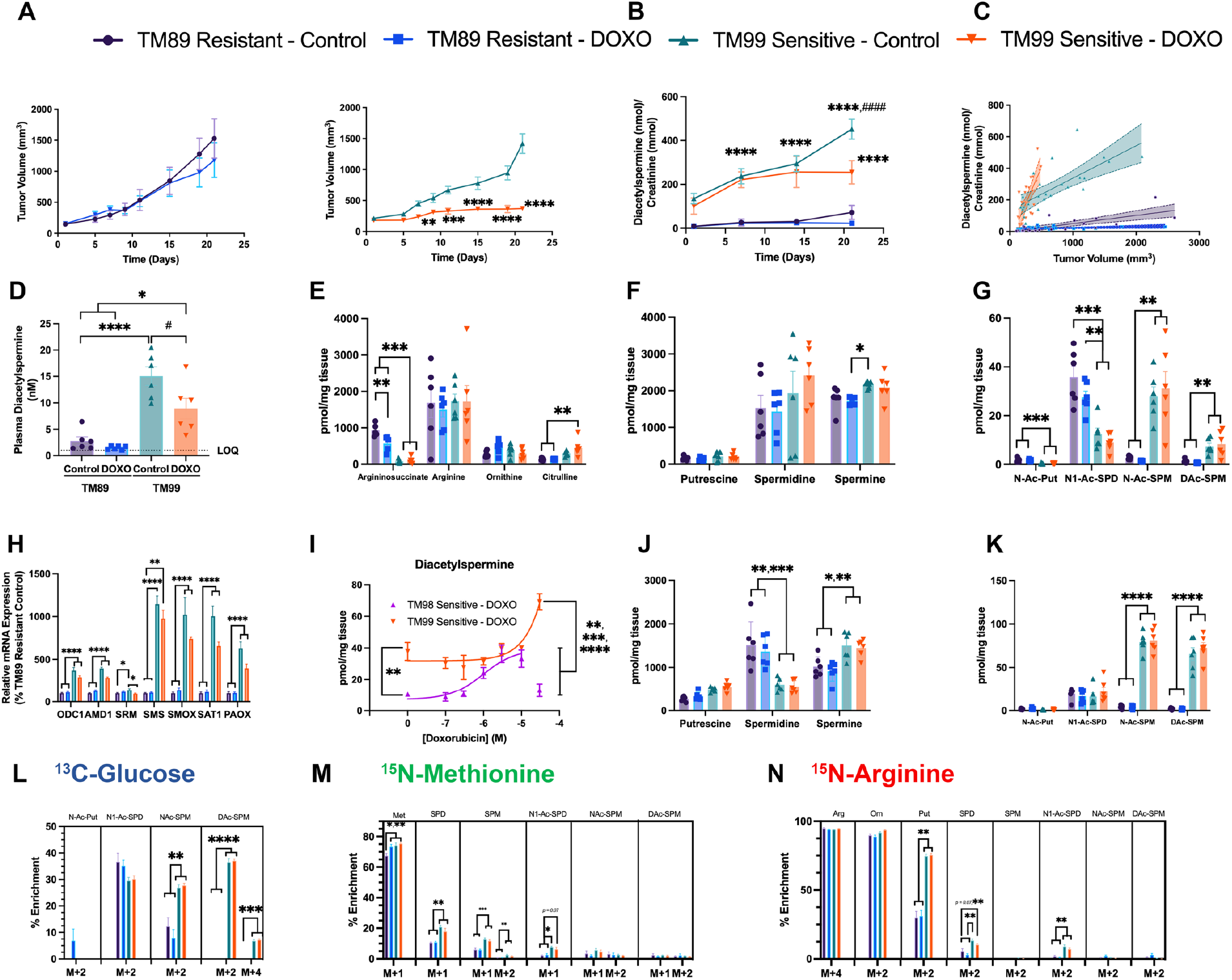
Effective doxorubicin treatment decreases urinary diacetylspermine in sensitive TNBC-PDX with high baseline SAT1 function. **(A**) Tumor growth curves showing TM89 resistant and TM99 sensitivity after treatment with vehicle (5% dextrose, IV) or doxorubicin (DOXO, 2 mg/kg, IV), mean ± s.e.m., n = 6. (**B**) Quantified urinary diacetylspermine and **(C)** correlation with tumor volume. **D**) Plasma diacetylspermine and tumor urea cycle (**E**), polyamine (**F**) and acetylated polyamine (**G**) metabolites from TM89 and TM99 TNBC-PDX mice, n = 6. (**H**) Doxorubicin dose-response in TM99 and TM98 *ex vivo* TNBC-PDX tissue slices, n=3. Urea cycle (**I**), polyamine (**J**) and acetylated polyamine (**K**) metabolites demonstrating elevated spermine and spermine acetylation in TM99 compared to TM89 *ex vivo* tissue slices, independent of doxorubicin (1μM) treatment. Stable isotope metabolic flux in TM89 and TM99 *ex vivo* tissue slices treated with ^13^C-glucose (**L**), ^15^N-methionine (**M**), and ^15^N-arginine (**O**), vehicle (DMSO) or 1 μM doxorubicin for 24 hours. *Ex vivo* slice experiments were performed in triplicate, n = 6. All data are presented as mean ± s.e.m., \**P*<0.05, \*\**P*<0.01, \*\*\**P*<0.01, \*\*\*\**P*<0.0001; ^#^ *P*<0.05, ^####^ *P*<0.0001 vs TM99 control. Significance was determined by two-sided one-way ANOVA with Tukey’s test or two-way repeated measures ANOVA with Sidak’s correction.

Interestingly, TM99 responded to doxorubicin treatment through a reduction in tumor size, but tumor diacetylspermine production was unaffected. Measuring *SAT1* mRNA revealed a striking 10-fold greater abudance in TM99 PDX expression compared to TM89 PDX, suggesting a high baseline level of SAT1 in TM99 (Figure 4H). Furthermore, in TM99 *ex vivo* tumor slices, baseline diacetylspermine levels were 3.7-fold higher than in TM98 (Figure 4L). When TM99 *ex vivo* slices were treated with doxorubicin, a concentration above physiological relevance was necessary to increase diacetylspermine beyond the already elevated baseline production. Therefore, TM99 tumor production of diacetylspermine was unaffected by doxorubicin in vivo but the resulting urine levels predictably decreased in proportion to decreasing tumor size. Indeed urine, plasma and tumor diacetylspermine significantly correlated across all TNBC-PDX models (Supplemental Figure 4).

In TM99, in vivo tumor tissue and *ex vivo* slices demonstrated elevated N-acetylspermine and diacetylspermine levels compared to TM89, but N1-acetyspermidine was either unaffected or decreased, respectively (Figure 4G and 4K). Unlike TM98, acetylated spermine metabolites were observed as the major catabolic product of high SAT1 baseline function in TM99 (Figure 4L and Supplemental Figure 5). This observation was further supported by increased ^13^C-enrichment of N-acetylspermine and diacetylspermine, but not N-acetylspermidine in TM99 (Figure 4L). Consistent with the TM98-doxorubicin sensitive tumor, ^15^N-derived from methionine demonstrated increased polyamine flux in TM99 compared to TM89 (Figure 4M). However, downstream ^15^N-methionine enrichment into acetylated spermine metabolites was undetected in TM99 *ex vivo* slices. Together these data suggest that in tumors with high baseline SAT1 function, spermine acetylation is the major route of SAT1 metabolism and elevated polyamine flux may not contribute to acetylated metabolites.

### Diacetylspermine is elevated in breast cancer tumors

Our studies in TNBC-PDX indicated that increased urine diacetylspermine is directly produced by the TNBC tumor. Therefore, to determine if diacetylspermine is altered in clinical breast cancer tumors, we measured urea cycle, polyamines and acetylated polyamines in breast cancer tumor samples and matched non-tumor tissue. Polyamines and acetylated polyamines, including diacetylspermine, were significantly increased in tumor tissue compared to matched non-tumors (Figure 5A). When evaluating breast cancer tumors by molecular subtype, HER2+ and TNBC tumors produced significantly higher levels of spermidine, spermine and acetylated polyamines compared to ER+ tumors (Figure 5E-F). Moreover, gene expression data from The Cancer Genome Atlas and METABRIC studies demonstrate that SAT1 expression is significantly increased in HER2+ and TNBC tumors compared to ER+ (Figure 5G, H).

**Figure 5.**
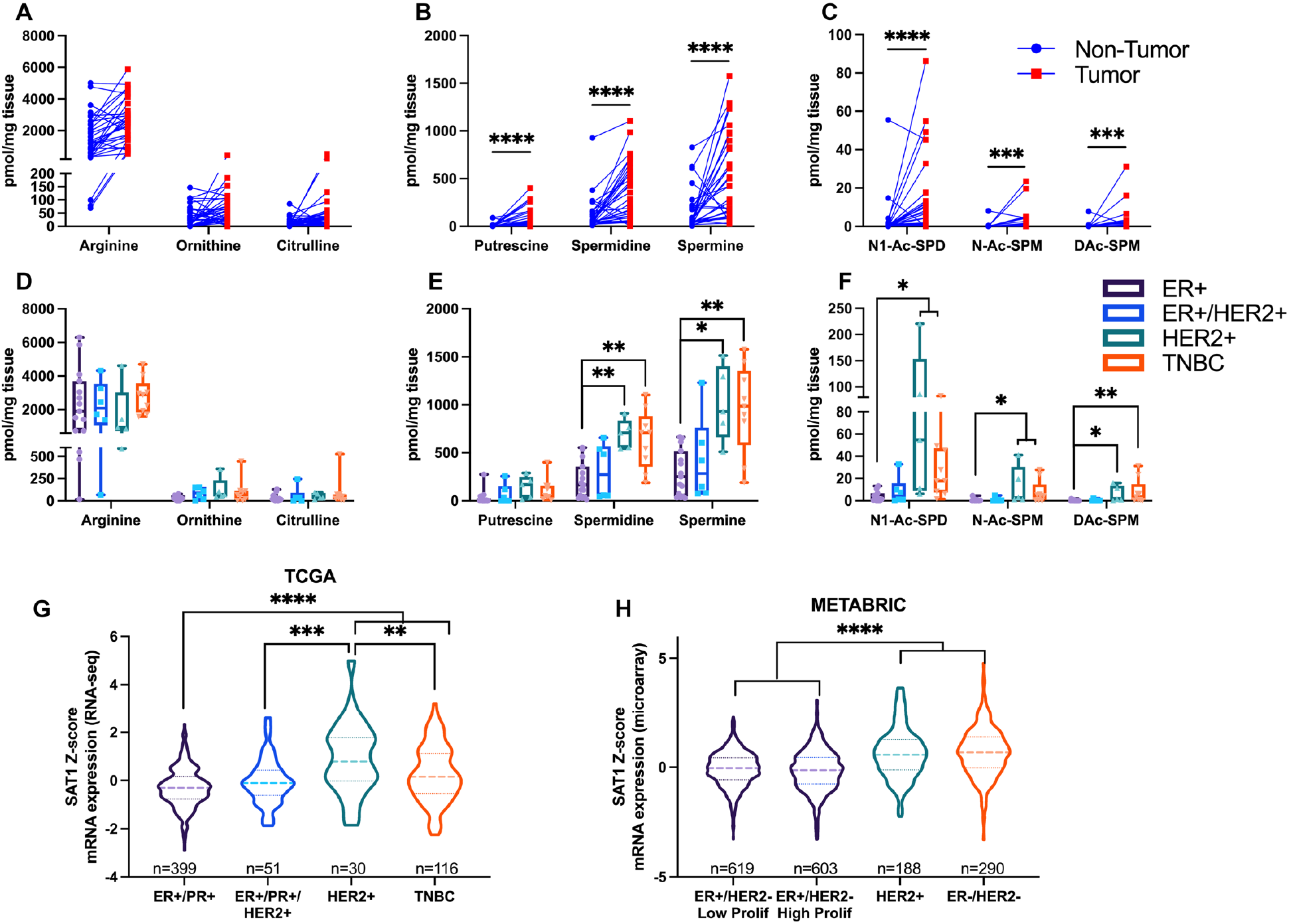
Diacetylspermine is elevated in breast cancer tissue, specifically in TNBC and HER2+ molecular subtypes. Breast cancer tumor vs matched non-tumor tissue levels of urea cycle (**A**), polyamine (**B**) and acetylated polyamine metabolites (**C**), n = 33. Urea cycle (**D**), polyamine (**E**) and acetylated polyamine (**F**) metabolites in breast cancer according to molecular subtype classification. (**G**) SAT1 mRNA expression in TCGA and **h**, METABRIC studies were obtained from cBioPortal. Box and swarm plots represent interquartile range (IQR) with center median and 1.5 x IQR whiskers. Gene expression data are presented in violin plots with n values indicated in the figure. Significance was determined by two-sided Wilcoxon matched-pa or one-way ANOVA with Tukey’s test.

### Urine diacetylspermine increases with tumor size and is elevated in patients with high grade tumors

Given that urine diacetylspermine correlated with TNBC-PDX tumor volume, we next asked whether urine levels were also associated with elevated tumor size in breast cancer patients from a large clinical study. A subset of 12-hour urine samples from the Polish Breast Cancer Study (n=717) containing control (n=200), ER+/PR+ (n=200), ER+/HER2 (n=36), ER-/PR-/HER2+(n=81), and TNBC (n=200) samples, including 500 breast cancer patients with tumor volume information, were analyzed (NCT00341458). Remarkably, urine diacetylspermine levels progressively increased with increasing tumor size in breast cancer patients (Figure 6A). In addition, urine diacetylspermine was significantly increased in patients with poorly differentiated tumors compared to lower grades (Figure 6B). However, there was no difference between urine diacetylspermine levels in control subjects and breast cancer patients (Figure 6C).

**Figure 6.**
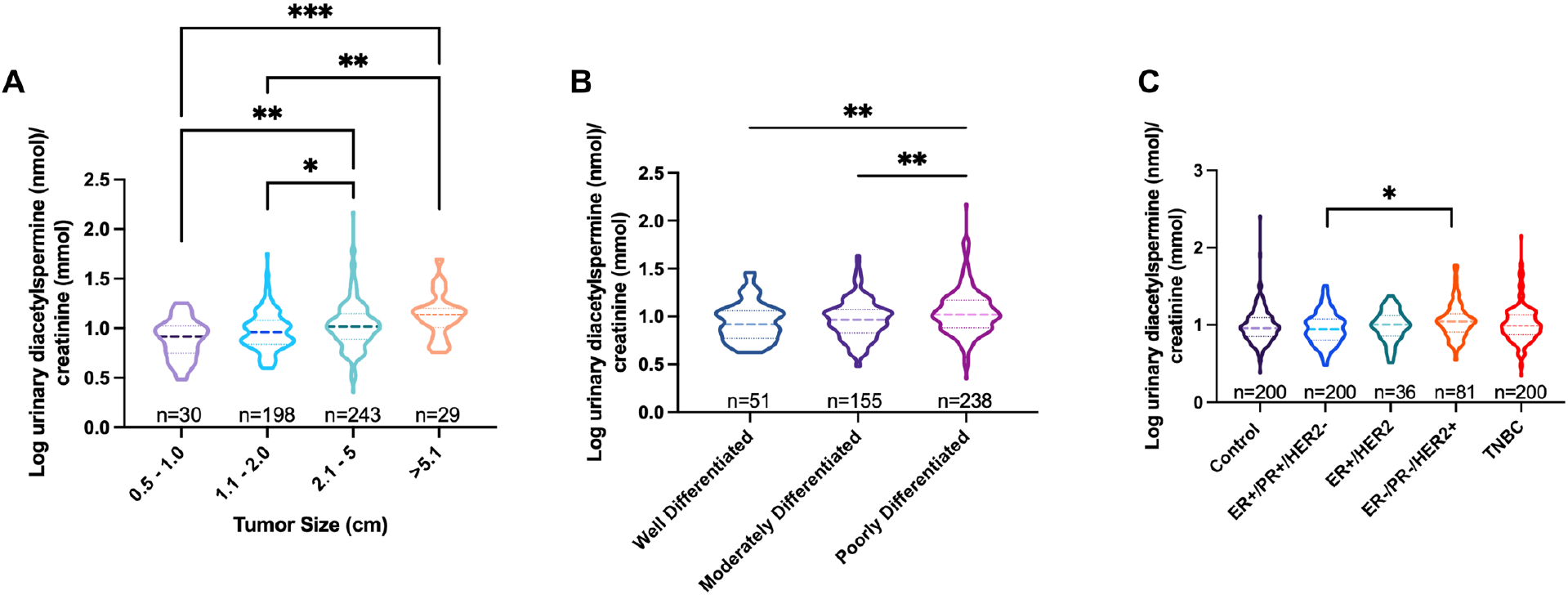
Urine diacetylspermine levels increase with tumor volume and are elevated in patients with poorly differentiated tumors. 12-hour urine diacetylspermine levels relative to tumor volume (**A**), tumor grade (**B**) and molecular subtype (**C**) in breast cancer patients from the Polish Breast Cancer Study (NCT00341458). Diacetylspermine concentrations are log transformed and normalized to creatinine. Data are presented as violin plots with n values indicated in the figure. Significance was determined by two-sided one-way ANOVA with Tukey’s test.

## Discussion

Tumor-derived metabolites excreted into blood and urine can be sampled non-invasively to provide an indication of treatment status. Moreover, when these metabolites are also directly altered by the treatment, they can act as an early signal of treatment effectiveness. Here, we conducted a highly controlled pharmacometabolomics study using PDX models and identified diacetylspermine as a metabolic biomarker of doxorubicin effectiveness in TNBC. We then perform a rigorous analysis to characterize tumor diacetylspermine production and determined its potential utility to assess tumor size by measuring urine levels in breast cancer patients.

Our aim was to design a preclinical study that would optimize biomarker identification and remain relevant to clinical sample collection. Therefore, longitudinal urine was collected from TNBC-PDX mice at the time of drug treatment (14). Repeated-measures analysis resulted in the identification of diacetylspermine as the most significantly altered metabolic feature in our urine metabolomics analysis. Interestingly, diacetylspermine was first discovered as a urine cancer biomarker in patients (15) and was identified in breast, colon and lung cancers, but with few supporting studies in animal cancer models (16-18). Therefore, the identification of a known cancer biomarker as the top candidate in TNBC-PDX models highlights the translational value of this approach.

Urine diacetylspermine clearly demonstrated a longitudinal correlation consistent with tumor growth in the TM98 and TM99 doxorubicin sensitive TNBC-PDX models. However, when TM98 mice were treated with doxorubicin and tumor volume declined, we found a further increase in urine diacetylspermine. Tumor measurements, *ex vivo* slices and *in vitro* experiments determined that SAT1 expression was directly induced by doxorubicin, thereby resulting in elevated urine diacetylspermine. This effect is consistent with data from the NCI-60 cell lines treated with doxorubicin, and dose-response curves were superimposable for breast cancer cell lines and TM98 *ex vivo* slices.

We collected urine from mice 7 days after the final doxorubicin dose and, in doing so, we discovered that urine diacetylspermine levels continued to increase long after doxorubicin had been systemically cleared. Indeed, the effects of doxorubicin on gene expression were shown to outlast its elimination (19). Upon further study, we demonstrated that elevated SAT1 occurs 24 hours after treatment and that diacetylspermine production lasts for 14 days following the final doxorubicin dose. These data suggest a clinically amenable timeframe for measuring diacetylspermine production as an indicator of doxorubicin effectiveness in TNBC.

Cancer cells require polyamines for growth and proliferation. To meet this demand, altered signaling pathways during cancer reprogramming upregulate rate limiting anabolic enzymes, ODC1 and AMD1 (20). Elevated SAT1 and decreased levels of PAOX were previously demonstrated in breast cancer, creating a favorable environment for diacetylspermine production (21). SAT1 responds to elevated free polyamines by increasing polyamine acetylation (22). This mechanism was shown in a breast cancer cell line (23) and was present when *ex vivo* slices were treated with spermine regardless of doxorubicin sensitivity or baseline SAT1 expression and diacetylspermine production. A significant advantage of TNBC-PDX models is the ability to resect tumors and culture *ex vivo* slices, thereby directly assessing tumor metabolic flux using stable isotope tracers (24, 25). Spermidine and spermine levels were similar between TM98 and TM97 in *ex vivo* slices; however, ODC1 and AMD1 expression as well as incorporation of ^15^N from methionine and arginine were elevated in TM98, demonstrating increased polyamine flux. Increased tracer levels were also evident in acetylated metabolites. These data suggest that acetylation may be increased to keep polyamine levels under tight control, thereby leading to elevated diacetylspermine. Conversely, high-baseline expression and activity of SAT1 in TM99 did not necessitate elevated polyamine flux for the generation of acetylated polyamines, suggesting an alternate mechanism of increased SAT1 expression at baseline in this model. Furthermore, the preferential generation of acetylated spermine metabolites in TM99 may be indicative of spermine availability (23).

Doxorubicin-mediated induction of SAT1 demonstrated a pattern similar to previous evidence of overexpression, whereby putrescine levels increase in response to polyamine depletion, further indicating a direct effect on SAT1 (26). A recent study described a doxorubicin-mediated decrease in ODC1 activity 48 hours after treatment, which suggests that the increase at 24 hours may dissipate (27). This effect may contribute to reduced tumor growth; however, ODC1 levels were unaltered 7 days after doxorubicin treatment in vivo. In addition, polyamine flux was elevated during SAT1 overexpression in a previous study but, in *ex vivo* tissue slices, doxorubicin treatment only affected acetylated polyamine flux (26). Together, these data confirm that doxorubicin treatment mainly affects diacetylspermine levels by modulating SAT1 expression and activity.

Although doxorubicin largely induces SAT1 expression at physiological concentrations as demonstrated in TM98 and NCI-60 cell lines, doxorubicin decreased plasma and urine diacetylspermine in TM99, which coincided with decreasing tumor size. However, urine and *ex vivo* tumor diacetylspermine concentrations as well as ^13^C-glucose tracer incorporation were greater than two-fold higher in TM99 than TM98. Our data provide evidence that the mechanism of doxorubicin induction of SAT1 did occur in TM99, but only at supraphysiologic concentrations. We propose that high-baseline levels of SAT1 and diacetylspermine production mask the induction by doxorubicin at physiological levels. Therefore, these data shed light on the potential use of urine diacetylspermine as an indicator of therapeutic effectiveness when the treatment does not directly affect SAT1 expression and function. Under these conditions, effective therapy is predictably indicated by a decrease in urine diacetylspermine.

The utility of diacetylspermine as a therapeutic biomarker depends on elevated tumor production. We compared polyamines and acetylated polyamines in tumor tissue with matched control tissue and found that all polyamines and acetylated polyamines were significantly elevated. Therefore, the majority of breast cancer tumors produce elevated levels. Notably, acetylated polyamines were increased in HER2+ and TNBC tumors when compared to ER+ molecular subtypes, which was consistent with TCGA and METABRIC expression levels of SAT1. Chemotherapy is commonly used to treat TNBC and HER2+ breast cancer (28) and therefore, these patients are more likely to benefit from therapeutic biomarker monitoring.

Diacetylspermine excreted from tumor tissue is readily cleared into urine creating a reservoir to measure in vivo production (29). Although several studies have identified diacetylspermine as a cancer biomarker, this is the first to demonstrate that urine diacetylspermine increases with increasing tumor volume in TNBC-PDX mice and in breast cancer patients. Moreover, elevated urine diacetylspermine was found in patients with poorly differentiated tumors, consistent with previous evidence in colorectal cancer (30). Others recently demonstrated elevated plasma diacetylspermine in TNBC patients when compared to healthy subjects (18). In contrast, we did not find significant differences in urine diacetylspermine between control and breast cancer patients. This may be explained by interindividual variability, and highlights the need for baseline and longitudinal sampling, similar to our TNBC-PDX study design. Importantly, these data suggest that breast cancer tumor size is associated with urine diacetylspermine, which is a fundamental feature for a biomarker of therapeutic effectiveness.

There are several limitations to our study. Although we identified doxorubicin sensitive TNBC-PDX models with elevated diacetylspermine production, consistent with human tumors, all doxorubicin-resistant models did not produce significant levels of diacetylspermine. Future studies are necessary to determine whether low diacetylspermine is a characteristic of doxorubicin resistance. Breast cancer tumor and urine samples were not obtained from the same study and therefore, we are unable to directly compare tumor and urine levels in patients. Only a single treatment naïve urine sample was obtained from breast cancer patients, therefore longitudinal sampling and response to doxorubicin treatment remains to be evaluated clinically.

Our study has several strengths. First, all TNBC-PDXs consistently demonstrated sensitivity or resistance to doxorubicin over multiple passages throughout the study, confirming the robustness of these models. Second, TNBC tumor variability was captured by using multiple TNBC-PDX models to provide a broad picture of diacetylspermine production and response to doxorubicin treatment. Third, culturing *ex vivo* tumor slices allowed us to assess polyamine metabolism and metabolic flux directly from our in vivo models. Finally, we quantified urine diacetylspermine from a large breast cancer case-control study that collected urine.

In conclusion, using a pharmacometabolomics approach, we discovered that urine diacetylspermine is a metabolic biomarker of doxorubicin effectiveness in TNBC-PDX models and is associated with tumor volume. Our findings provide evidence that urinary diacetylspermine may be a clinical biomarker of breast cancer treatment effectiveness. Moreover, this approach can be applied to other cancer types and drug treatment regimens. Future prospective clinical studies are necessary to determine the utility of urinary diacetylspermine as a guide to precision therapy.

## Methods

### Mouse models and cell lines

Triple negative breast cancer patient derived xenograft models TM00089 (TM89 doxorubicin-resistant, BR0620F), TM00097 (TM97 doxorubicin-resistant, BR1077F), TM00098 (TM98 doxorubicin-sensitive, BR1126F) and TM00099 (TM99 doxorubicin-sensitive, BR1367F) were obtained from Jackson Laboratories between passages 3 and 7. All animal studies were approved by the NCI Institutional Animal Care and Use Committee (IACUC). NSG mice were obtained for NCI Frederick or Jackson Laboratories. BT549 and HS578T cells were obtained from the NCI Developmental Therapeutics Program (DTP) repository.

### TNBC-PDX in vivo studies

TNBC-PDX tumors were surgically resected from TNBC-PDX mice, cut into 2 × 2 × 2 mm and subcutaneously implanted into 5-8-week-old female and mice. When tumors reached 100-200 mm^3^, TNBC-PDX mice were randomized and placed into control or doxorubicin groups. Subsequently, mice were given doxorubicin (2 mg/kg, Caymen Chemicals) or vehicle (5% dextrose) by intravenous injection into the lateral tail vein once weekly for 3 weeks. Tumors were measured thrice weekly using calipers and tumor volumes were calculated using the following formula: volume = (length x width^2^)/2. For urine collection, mice were placed in autoclaved glass metabolic cages (Jencons, Metabowl) and contained in a HEPA-filtered laminar flow hood for 24 hours. TNBC-PDX mice were placed in metabolic cages on days 1, 7 and 14, immediately following each drug administration. At 21 days, a final 24-hour urine collection was performed in the absence of drug treatment followed by euthanization to collect plasma and tissue, unless otherwise indicated. Urine samples were centrifuged at 2000 g following and the supernatant was frozen at -80 °C.

### Ex vivo tissue slice experiments

Untreated TNBC-PDX mice with tumors between 1000 and 1500 mm^3^ were euthanized and tumors excised under sterile conditions. Tumors were sliced once along the major-axis to create a flat surface and covered with 2% low melting-point agarose heated to ∼40 °C (25). When agarose was hardened, tumors were cut into 500 μm slices using a McIlwain Tissue Chopper (Ted Pella, Inc.) and washed twice in PBS. *Ex vivo* slices were placed in 6-well plates containing 3 mL of media then onto an orbital shaker in a humidified 37 °C incubator under 5% CO_2_. RPMI containing 10 % FBS, 100 U/mL penicillin/streptomycin/antimycotic was used to culture *ex vivo* slices. For isotopic labelling experiments, tumor slices were cultured in RPMI containing 10% dialyzed FBS, 100 U/mL penicillin/streptomycin/antimycotic and unlabeled or uniform labelled ^13^C-glucose (in RPMI cat. 11879020), ^15^N-methionine (in RPMI cat. A1451701) or ^15^N-arginine (in RPMI cat. 88365 plus L-lysine). After 24 hours, tissue slices were removed, washed with PBS, weighed, blotted dry and flash frozen in liquid nitrogen. Viability after 24 hour treatment was compared to fresh *ex vivo* slices using PrestoBlue (cat. A13261), as previously described (25).

### Untargeted urine metabolomics analysis

Urine creatinine concentration was measured using the Jaffe method. To reduce urine concentration variability for metabolomics analysis, all samples were diluted to one standard deviation below the mean creatinine concentration. Samples were then diluted 1:5 in acetonitrile/water/methanol (65/30/5) containing 10 μM α-aminopimelic acid as internal standard and injected onto a Waters H-Class ultra-performance liquid chromatography (UPLC) coupled to a Waters Xevo G2 quadrupole time of flight MS (QTOFMS) in positive and negative ionization mode, as previously described (31). Urine metabolomics raw data files were processed using Progenesis QI software and normalized to creatinine. The resulting feature lists for positive and negative mode were combined after filtering features that ionize in both modes as previously described (32).

### Polyamine and Amino acids targeted LC-MS analysis

Urine and plasma samples were diluted 1:3 with 6% trichloroacetic acid (TCA) (33) containing internal standards (1 μM 1,7-diaminoheptane, 100 nM d6-diacetlyspermine and 10 μM d8-spermine). Tissue and *ex vivo* slice samples were diluted with 20 μL/mg of tissue in 6% TCA containing internal standards and processed in a Precellys homogenizer (6500 RPM for 30s x 2). Samples were centrifuged at 15,000 g for 10 min and 10 μL of supernatant was added to 70 μL of 100 mM borate buffer containing 100 μM NaOH (check pH in lab book), 1 mM ascorbic acid and 1 mM TCEP (34). Samples were derivatized with 20 μL of 6-aminoquinolyl-N-hydroxysuccinimidyl carbamate (AQC, AccQ-Tag, Waters) followed by a 10-minute incubation at room temperature and a 10-minute incubation at 55 °C to quench the reaction. The resulting derivatized sample was injected onto a Waters Acquity BEH Phenyl-Hexyl column (2.1 × 50 mm) maintained at 40 °C and a flow rate of 0.6 mL/min in a Waters I-Class UPLC. The mobile phase consisted of water + 0.1% formic acid (A) and acetonitrile + 0.1% formic acid (B) and run under the following conditions: 0-0.5 mins, 0 %B; 0.5-4 mins, 0-15% B; 4-6 mins 15-30% B; 6-6.5 mins, 30-99% B, 7.5-9 mins 0% B. Eluting metabolites were measured in a Waters Xevo TQS using multiple reaction monitoring (MRM, Supplemental Table 2). Parent m/z values were calculated using the following equation:

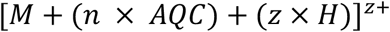

where *n* is the number of derivatized amines with *AQC* and *z* is the charge number of the ion. The most sensitive ion charge was used for each metabolite. Spermine was normalized to d8-spermine and all other metabolites were normalized to d6-diacetylspermine or diluted 20-fold and normalized to 1,7-diaminoheptane.

### Stable isotope metabolic flux analysis

*Ex vivo* slices incubated with media containing stable isotope labelled compounds were processed as described above and measured using a Waters Synapt G2-S operating in positive mode with a 0.5 kV and 40 V, capillary and cone voltage, respectively. The source temperature was set to 150 °C and the desolvation gas flow rate was 950 L/hour at 500 °C. Ions generated from isotopic enrichment of ^13^C or ^15^N were monitored for amino acids and polyamines as described in Supplemental Table 2. IsoCorrectoR was used to correct for natural isotope abundance (35).

### Human specific primer design and qPCR

Total RNA was extracted using Trizol and cDNA synthesized with qScript cDNA Supermix, following manufacturers protocols. Relative mRNA quantified with PerfeCTa SYBR Green (Quantabio) and gene expression normalize to human GAPDH using the delta-delta *C*_*T*_ method. Human specific primers were designed in non-overlapping regions of mouse and human transcript orthologs. Human primer specificity was validated by qPCR using cDNA generated from BT-549 human breast cancer cells and mouse liver tissue.

### Human tissue and urine specimens

Fresh-frozen tumor samples were obtained from unselected breast cancer patients at all disease stages having a mastectomy between February 15, 1993, and August 27, 2003, under the NCI resource contract “Collection and Evaluation of Human Tissues and Cells from Donors with an Epidemiology Profile”, as previously described (36). All patients provided written informed consent. Urine samples were obtained from the Polish Breast Cancer Study, which comprised of a population-based breast cancer case-control study in Poland between 2000 and 2003, as previously described (37) (NCT00341458). Urine samples were randomly selected from the 2241 control and 1962 breast cancer study subjects as follows: 200 control, ER+/PR+, and TNBC as well as all ER+/PR+/HER2+ (n=31) and HER2+ (n=81).

### Analysis of public data

Data were downloaded from The NCI Transcriptional Pharmacodynamics Workbench were downloaded from GSE116436 (38). SAT1 expression data from METABRIC and TCGA were downloaded from cBioPortal (39, 40).

### Statistics

Principal component analysis was performed on resulting features in SIMCA (version 15). For univariate statistical analysis, features were analyzed by repeated measure two-way ANOVA using Metaboanalyst (41). P values were adjusted according to the Benjamini Hochberg procedure and q < 0.05 was considered significantly different. All other statistical analysis methods were performed as indicated in figure legends using GraphPad Prism version 8 and 9.

## Author contributions

Conceptualization: TJV; Data curation: TJV; Formal analysis: TJV, ST; Funding acquisition: FJG;Investigation: TJV, ST, KWK, KH; Methodology: TJV, KWK, ST; Supervision: FJG, ST, SA; Writing original draft: TJV; Writing – review & editing: all authors

## Acknowledgements

We wish to acknowledge John Buckley for technical assistance The study was funded by the National Cancer Institute Intramural Research Program. TJV was supported by a Canadian Institutes of Health Research (CIHR) Postdoctoral Fellowship. The Polish Breast Cancer Study was funded by the Intramural Research Program in the Division of Cancer Epidemiology and Genetics, the US National Institutes of Health (NIH), National Cancer Institute (NCI). The authors acknowledge Dr. Montserrat Garcia-Closas and Dr. Thomas U. Ahearn for their assistance in providing access to data and samples from the Polish Breast Cancer Study. We thank the physicians, pathologists, nurses, interviewers, and study participants that contributed to the success of the Polish Breast Cancer Study.

