## Supplemental Information for "Pharmacometabolomics reveals urinary diacetylspermine as a biomarker of doxorubicin effectiveness in triple negative breast cancer"

Supplementary Figures

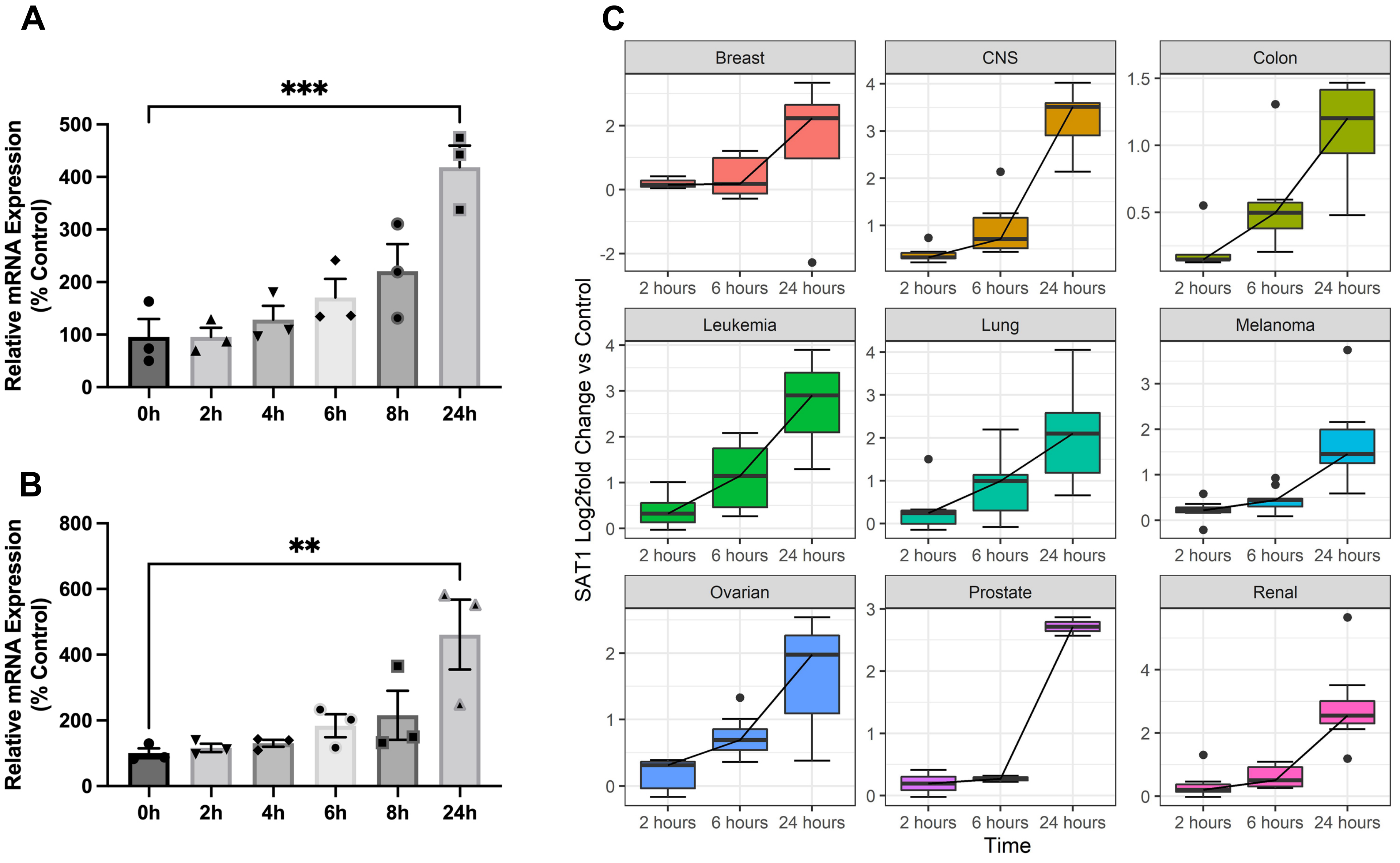


**Supplemental Figure 1. SAT1 induction by doxorubicin is time-dependent.** SAT1 mRNA expression after 1 µM doxorubicin treatment at the indicated time points in BT549 (**A**), HS578T (**B**) and NCI-60 cells lines (**C**). Data are presented as mean s.e.m , n = 3 (A,B), or were downloaded from GSE116436 and are presented as box and whisker plots of SAT1 $log_2$-fold change compared to control at the same timepoint (C) (Monks et al., 2018). Significance was determined by one-way ANOVA with Tukey's test.

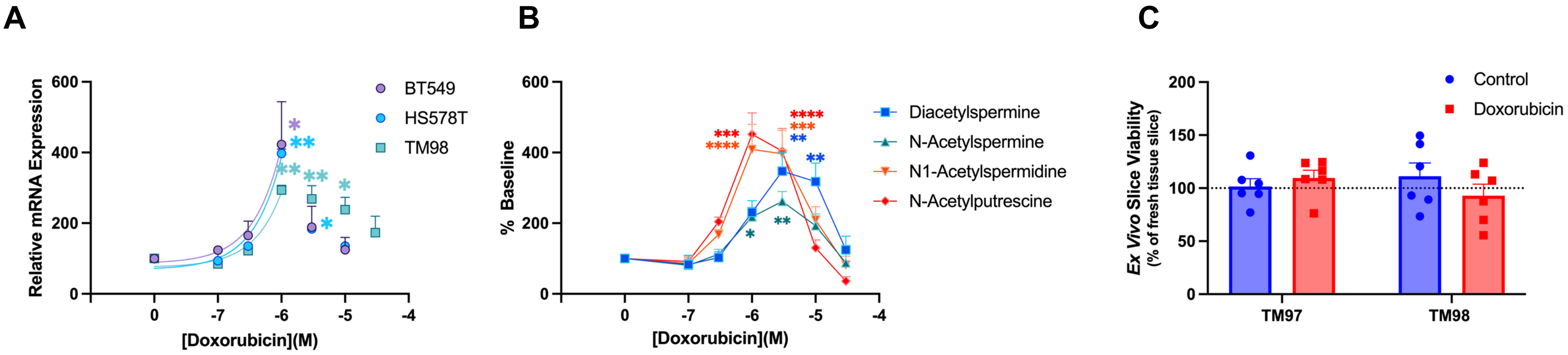


**Supplemental Figure 2. Doxorubicin dose-dependently induces SAT1 mRNA expression and polyamine acetylation.** BT549, HS578T and TM98 SAT1 mRNA expression (**A**) and polyamine acetylation (**B**) 24 hours after treatment with doxorubicin (n=3). **C**, Tissue viability of TM97 and TM98 ex vivo slices 24 hours after treatment with vehicle or doxorubicin and normalized to fresh ex vivo slices (n=6). Experiments were performed in triplicate. Data are presented as mean ± s.e.m. *$P$<0.05, **$P$<0.01, ***$P$<0.001, ****$P$<0.0001 compared to control. Significance was determined by one-way ANOVA with Tukey's test.

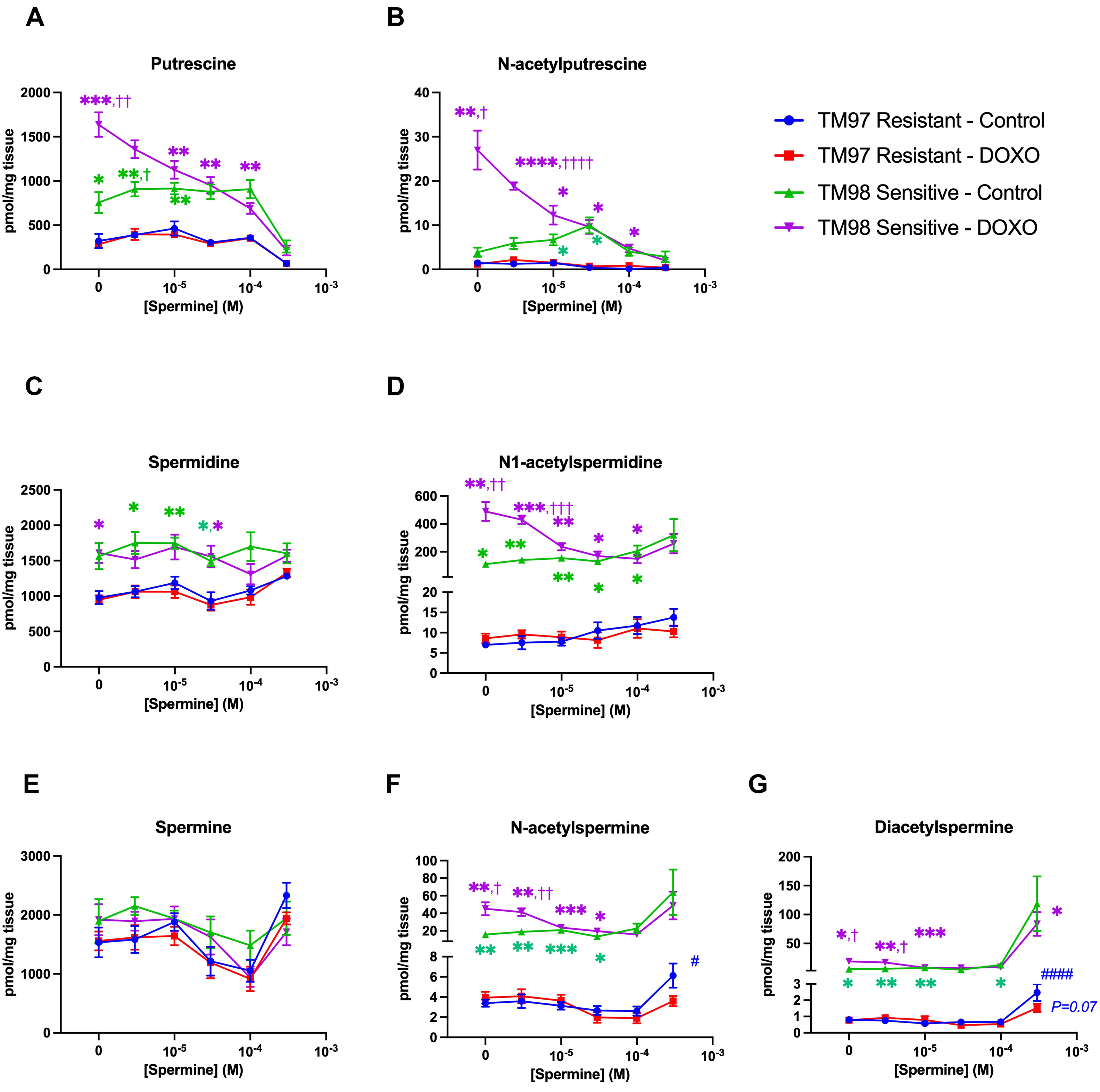


**Supplemental Figure 3. Spermine dose-dependently reverses doxorubicin induced polyamine acetylation in TM98, but at high concentrations, spermine increases diacetylspermine production regardless of doxorubicin sensitivity.** Putrescine (**A**), N-acetylputrescine (**B**), spermidine (**C**), N1-acetylspermdine (**D**), spermine (**E**), N-acetylspermine (**F**) and diacetylspermine (**G**) in TM97 and TM98 *ex vivo* tissue slices treated with vehicle or doxorubicin (DOXO) 1 μM in the presence of spermine for 24 hours. Experiments were performed in duplicate and data are presented as mean ± s.e.m, n = 6. **P*<0.05, ***P*<0.01, ****P*<0.001, *****P*<0.0001 vs TM97; $^{\#}$ *P*<0.05, $^{\#\#\#\#}$ *P*<0.0001 vs TM97 in the absence of spermine; $^{\dagger}$*P*<0.05, $^{\dagger\dagger}$*P*<0.01, $^{\dagger\dagger\dagger}$*P*<0.001, $^{\dagger\dagger\dagger\dagger}$*P*<0.0001 vs TM98 control. Significance was determined by one-way ANOVA with Tukey's test.

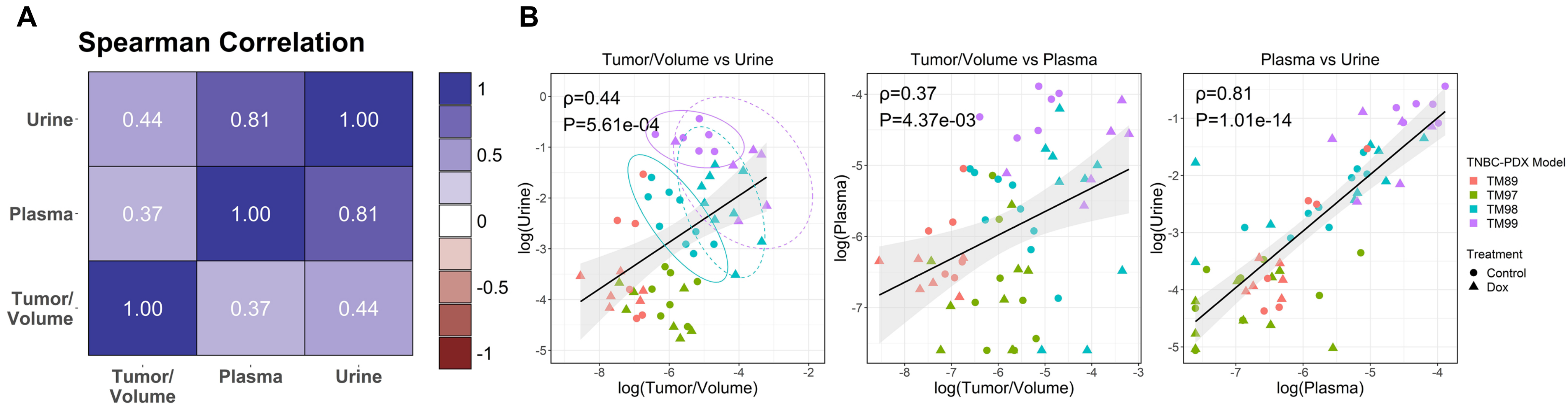


**Supplemental Figure 4. Urine diacetylspermine directly correlates with plasma and tumor levels.** (**A**), Spearman rank correlation coefficients between urine, plasma and tumor/volume diacetylspermine levels. (**B**), individual correlation plots show that untreated TM99 produces more urine diacetylspermine per tumor volume than other tumors and that doxorubicin treatment increases TM98 diacetylspermine tumor production, which is reflected in urine. Tumor concentration was normalized to tumor volume and urine diacetylspermine was normalized to creatinine. All data were log transformed; TM97 and TM98 n = 8-9, TM89 and TM99 n = 6. Significance was determined by spearman correlation and P values are annotated.

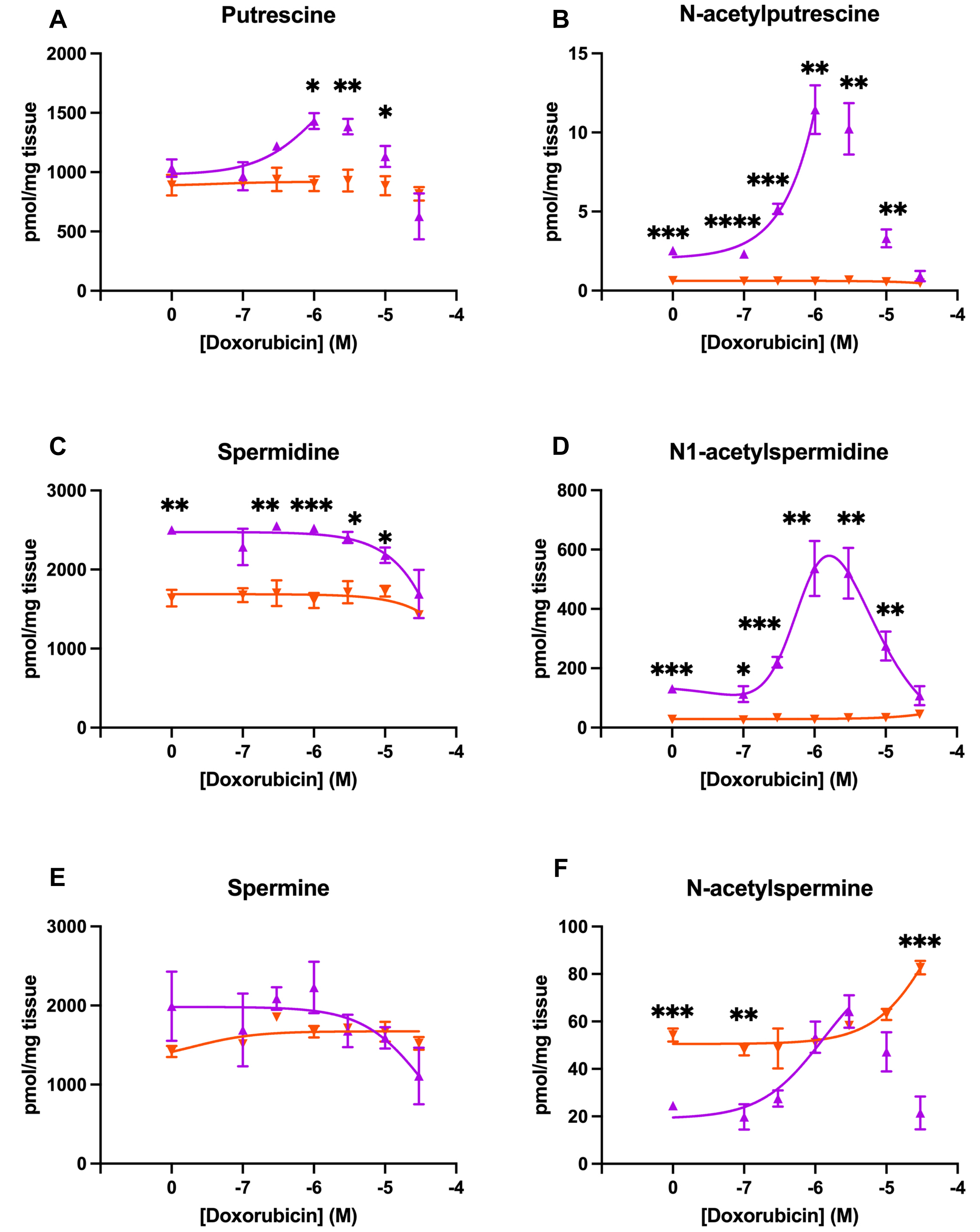


**Supplemental Figure 5. Doxorubicin dose-dependently induces polyamine acetylation in TNBC-PDX with low baseline SAT1 function, but preferentially increases spermine acetylation in TNBC-PDX high baseline SAT1 function.** Putrescine (**A**), N-acetylputrescine (**B**) spermidine(**C**), N-acetylspermidine (**D**), spermine (**E**) and N-acetylspermine (**F**) in TM98 (▲) and TM99 (▼) ex vivo tissue slices 24 hours after treatment with doxorubicin. Experiments were performed in triplicate and data are presented as mean ± s.e.m, n = 6. *$P$<0.05, **$P$<0.01, ***$P$<0.001, ****$P$<0.0001. Significance was determined by t-test. Data are fitted with three parameter or bell-shaped dose response curves.

**Supplemental Table S2.** List of AQC-derivatized MRM transitions and accurate m/z calculations for neutral and stable-isotope labeled urea cycle and polyamine metabolites

| | | | | | | | | | | 15N-labelling | | | | 13C-labelling | | | |
|---|---|---|---|---|---|---|---|---|---|---|---|---|---|---|---|---|---|
| Number | Name | Elemental Comp | M | n | Aqc(n) | H(n) | (n)+ | MRM transition | Accurate m/z | M+1 | M+2 | M+3 | M+4 | M+1 | M+2 | M+3 | M+4 |
| 1 | L-Arginine | C6H14N4O2 | 174.1117 | 1 | 170.048 | 1.007 | | 345.2 > 70 | 345.1667 | 346.1970 | 347.1941 | 348.1911 | 349.1881 | | | | |
| 2 | L-Argininosuccinic Acid | C10H18N4O6 | 290.1226 | 1 | 170.048 | 1.007 | 1 | 461.2 > 70 | 461.1776 | | | | | | | | |
| 3 | L-Citrulline | C6H13N3O3 | 175.0957 | 1 | 170.048 | 1.007 | 1 | 346.2 > 145 | 346.1507 | | | | | | | | |
| 4 | L-Ornithine | C5H12N2O2 | 132.0899 | 2 | 170.048 | 1.007 | 2 | 237.1 > 171 | 237.1000 | 237.5985 | 238.0970 | 238.5956 | 239.0941 | | | | |
| 5 | Putrescine | C4H12N2 | 88.1515 | 2 | 170.048 | 1.007 | 2 | 215.1 > 171 | 215.1308 | 215.6293 | 216.1278 | 216.6264 | 217.1249 | | | | |
| 6 | Spermidine | C7H19N3 | 145.1579 | 3 | 170.048 | 1.007 | 2 | 219.4 > 171 | 328.6580 | 329.1565 | 329.6550 | 330.1536 | 330.6521 | | | | |
| 7 | Spermine | C10H26N4 | 202.2157 | 4 | 170.048 | 1.007 | 2 | 221.6 > 171 | 442.2109 | 442.7094 | 443.2079 | 443.7065 | 444.2050 | | | | |
| 8 | N-Acetylputrescine | C9H21N3O | 130.1106 | 1 | 170.048 | 1.007 | 1 | 301.2 > 145 | 301.1656 | 302.1626 | 303.1597 | 304.1567 | 305.1537 | 302.1690 | 303.1723 | 304.1757 | 305.1790 |
| 9 | N1-Acetylspermidine | C9H21N3O | 187.1685 | 2 | 170.048 | 1.007 | 2 | 264.6 > 100 | 264.6393 | 265.1378 | 265.6363 | 266.1349 | 266.6334 | 265.1410 | 265.6427 | 266.1443 | 266.6460 |
| 10 | N-Acetylspermine | C12H28N4O | 244.2263 | 3 | 170.048 | 1.007 | 3 | 252.5 > 171 | 252.4638 | 252.7961 | 253.1285 | 253.4608 | 253.7932 | 252.7983 | 253.1327 | 253.4672 | 253.8016 |
| 11 | N1,N12-Diacetylspermine | C14H30N4O2 | 286.2369 | 2 | 170.048 | 1.007 | 1 | 314.2 > 100 | 627.3399 | 628.3369 | 629.3340 | 630.3310 | 631.3280 | 628.3433 | 629.3466 | 630.3500 | 631.3533 |
| 12 | d6-Diacetylspermine | C14H24D6N4O2 | 292.2745 | 2 | 170.048 | 1.007 | 1 | 317.2 > 171 | 633.3775 | | | | | | | | |
| 13 | d8-Spermine | C10H18D8N4 | 210.2660 | 4 | 170.048 | 1.007 | 2 | 446.2 > 171 | 446.2360 | | | | | | | | |
| 14 | 1,7-Diaminoheptane | C7H18N2 | 130.1470 | 2 | 170.048 | 1.007 | 2 | 236.1 > 171 | 236.1285 | | | | | | | | |
